# Characterisation of the immune repertoire of a humanised transgenic mouse through immunophenotyping and high-throughput sequencing

**DOI:** 10.1101/2022.06.27.497709

**Authors:** E Richardson, Š Binter, M Kosmac, M Ghraichy, V von Niederhausern, A Kovaltsuk, J Galson, J Trück, DF Kelly, CM Deane, P Kellam, SJ Watson

**Affiliations:** Kymab, a Sanofi Company, Babraham Research Campus, Cambridge, UK; Department of Statistics, University of Oxford, Oxford, UK; Division of Immunology, University Children’s Hospital, University of Zurich, Zurich, Switzerland; Children’s Research Center, University of Zurich, Zurich, Switzerland; Alchemab Therapeutics Ltd, Kings Cross, London, UK; Department of Paediatrics, University of Oxford, Oxford, UK; Department of Infectious Diseases, Faculty of Medicine, Imperial College London, UK

**Author notes:** Corresponding author: Simon Watson. Contributed equally to the study (E Richardson, Š Binter).

## Abstract

Immunoglobulin loci-transgenic animals are widely used in antibody discovery and increasingly in vaccine response modelling. In this study, we phenotypically characterised B-cell populations from the Intelliselect® Transgenic mouse (Kymouse) demonstrating full B-cell development competence. Comparison of the naïve B-cell receptor (BCR) repertoires of Kymice BCRs naïve human and murine BCR repertoires revealed key differences in germline gene usage and junctional diversification. These differences result in Kymice having CDRH3 length and diversity intermediate between mice and humans. To compare the structural space explored by CDRH3s in each species repertoire, we used computational structure prediction to show that Kymouse naïve BCR repertoires are more human-like than mouse-like in their predicted distribution of CDRH3 shape. Our combined sequence and structural analysis indicates that the naïve Kymouse BCR repertoire is diverse with key similarities to human repertoires, while immunophenotyping confirms that selected naïve B-cells are able to go through complete development.

## Introduction

Twenty-five years of progress in genetic engineering since the first Ig transgenic mouse (M. Brüggemann et al. 1989) culminated in 2014 in the insertion of the full set of variable human Ig genes in mice (Lee et al. 2014). Humanised immunoglobulin (Ig) loci-transgenic animal models have proven extremely useful in therapeutic antibody discovery and, more recently, in vaccine response modelling; twenty of the 127 therapeutic antibodies licensed in the US or EU as of April 2022 were derived from transgenic mouse platforms (data from Thera-SAbDab (Raybould et al. 2020)). Transgenic platforms have also found a new application in vaccine response modelling (Sok et al. 2016; Pantophlet et al. 2017; Walls et al. 2020). As humanised animal models become the source of a growing number of therapeutics and play an increasingly important role in the evaluation of novel vaccine candidates, it is crucial to understand the degree to which their B-cell repertoires can be considered representative of humans.

Contemporary Ig transgenic animal models vary according to the number of genes and localisation of the inserted human Ig loci (Green 2014; Brüggemann et al. 2015). In Kymab’s Intelliselect® Transgenic mouse (Kymouse), a complete set of human variable (V), diversity (D) and junction (J) genes of the IGH locus as well as the V and J genes of the IGλ and IGκ loci were inserted at the sites of the endogenous mouse loci. The mouse constant regions were retained, preserving downstream interactions with endogenous intracellular signalling components and cell membrane Fc receptors, resulting in functional, fully active chimeric antibodies. Kymice exhibit normal B-cell production and maturation and the resulting B-cell receptors (BCRs) are diverse, with human-like CDRH3 lengths and evidence of somatic hypermutation (Lee et al. 2014). However, the baseline phenotypic diversity in B-cells and B-cell receptors (BCRs) in the Kymouse has not been fully described.

B cells are an integral part of the humoral immune response due to their ability to produce antibodies against diverse antigens, providing protection against infection. They originate from hematopoietic stem cells in the bone marrow, where they undergo several phases of antigen-independent development leading to the generation of immature B cells. B cells are routinely classified based on their maturation status, antibody isotype, and effector function. Ig gene rearrangement during these early stages of B cell development results in the expression of a mature B cell receptor that is capable of binding to antigen. This is followed by positive and negative selection processes, which are designed to eliminate non-functional and self-reactive immature B cells. Surviving B cells complete antigen-independent maturation in the spleen, producing immunocompetent naïve mature B cells that subsequently develop into either follicular or marginal zone B cells. In response to vaccination or invading microbes, antigen-specific B cells within secondary lymphoid organs differentiate into antibody-producing cells, early memory cells, or rapidly proliferate and form structures known as germinal centres (Allen, Okada, and Cyster 2007). Germinal centres (GCs) are inducible lymphoid microenvironments that support the generation of affinity-matured, isotype-switched memory B cells and antibody secreting plasma cells. Long-lived plasma cells (PCs) secrete high-affinity antibodies, and memory cells can readily elicit an efficient antibody immune response upon re-exposure to the immune stimuli (Corcoran and Tarlinton 2016; Weisel and Shlomchik 2017). Iterative cycles of B cell hypermutation and selection within the GC leads to an accumulation of affinity-enhancing mutations and ultimately to the progressive increase of serum antibody affinity, a process known as antibody affinity maturation (Jacob et al. 1991). Antibody-secreting plasma cells play critical roles in protective immunity on the one hand and antibody mediated autoimmune disease on the other. During immune responses a small fraction of newly generated plasma cells enter either the bone marrow (BM) or the lamina propria of the small intestine where they populate specialised survival niches and become long-lived plasma cells (Lemke et al. 2016) thus maintaining antibody titres for extended periods.

The variable domain of a BCR is composed of a heavy chain and a light chain. Each of the chains in the antibody has three hypervariable regions known as the complementarity-determining regions (CDRs), which make most contacts with the antigen. The heavy chain locus consists of variable (V), diversity (D) and joining (J) gene segments, which recombine to form the heavy chain: the first two CDRs of the heavy chain, CDRH1 and CDRH2, are encoded by the V gene alone, while the third and most variable CDR, CDRH3, spans the V, D and J gene junctions. The insertion of random and palindromic nucleotides at the V-D and D-J junctions further contribute to the diversity of the CDRH3, ensuring binding diversity to different antigens and epitopes (Xu and Davis 2000). Each of the light chain loci, kappa and lambda, consist of V and J gene segments but no D gene segments, and both the germline as well as the recombined chain are less diverse than their heavy chain counterparts (Collins and Watson 2018).

Due to the greater diversity of the heavy chain, most next-generation sequencing of BCR repertoires (BCR-seq) has focused on the heavy chain; lower throughput methods exist for identifying the light chain pairing (Curtis and Lee 2020). The resulting BCR sequences can be aligned to reference germline gene databases to infer most likely germline gene origins and insertions or deletions at the V(D) or (D)J junctions (Ye et al. 2013). Alignment of BCRs to common germline genes also allows inference about clonal structure, as sequences sharing common germline gene assignments as well as homology in the CDRH3 loop may be inferred to have arisen from a common progenitor B-cell (Greiff et al. 2015; Yaari and Kleinstein 2015). The amino acid sequences of these heavy chains can also be functionally examined through annotation with structural tools (Kovaltsuk et al. 2017; Marks and Deane 2020). Changes in the pattern of CDRH3 shapes in BCR repertoires have been observed along the B-cell differentiation axis in both humans and mice (Kovaltsuk et al. 2020) but the extent to which the CDRH3 protein shape differs between humans and mice has not been explored.

Here we have characterised the frequency of germinal centre B cells, memory B cells and long-lived plasma cells from spleens, lymph nodes, and bone marrows of antigen-inexperienced Kymice (Lee et al. 2014). The frequencies of these B cell subsets as well as the breadth and nature of their BCR repertoires constitutes the first step in our understanding of how the immune system of this model organism responds to different antigens, vaccines, and pathogens that are both of scientific as well as therapeutic interest. Examining the nature of the naïve BCR repertoire in Kymice through both single-cell and bulk sequencing and structural analysis shows the Kymouse naïve BCR repertoires are more human-like in their distribution of CDRH3 shapes.

## Materials & Methods

### Kymouse data

Spleens, lymph nodes and bone marrows were collected from *n=*20 antigen inexperienced Intelliselect® Transgenic mice (Kymice). These Kymice contain chimeric immunoglobulin loci, with humanised variable domains (V_H_, V_K_, and V_L_) and a humanised lambda constant domain (C_L_), but murine heavy (C_H_) and kappa (C_K_) constant domains (Lee et al. 2014). The Kymice were selected to reduce any possible confounding effects by ensuring that: (i) there was an equal representation of sexes, (ii) mice were a range of ages on culling (6-12 weeks old), and (iii) mice were selected from different litters and culled over a period of 6 months.

### Lymphoid Cell Isolation and Cryopreservation

Bone marrow isolated from the femurs and tibias of each Kymouse were processed to single-cell suspensions by flushing the tissues with ice-cold FBS buffer and passing through 40 µm cell strainers. Spleens and inguinal lymph nodes were processed to single cell suspensions by homogenising through 40 µm cell strainers with ice-cold FBS buffer and pooled prior to staining and cell sorting. All single-cell suspensions were pelleted at 400 g for 10 minutes at 4°C prior to cryopreservation in 10% DMSO/FBS and storage in liquid nitrogen.

### Next Generation Sequencing (NGS) Analysis of Paired V_H_ and V_L_ Sequences from Single-Cell Sorted B Cells Derived from Kymice

For each Kymouse, the spleen and inguinal lymph nodes were processed to single-cell suspensions as described above before fluorescently-activated cell sorting (FACS) to recover CD19^+^ B220^+^ B cells into individual wells of a 96 well plate. RT-PCR was performed to amplify the V_H_ and V_L_ domains, and standard Illumina libraries were generated before sequencing on an Illumina MiSeq sequencer. The Change-O pipeline (Gupta et al. 2015) was used to process the sequence data; naïve BCR sequences were characterized as immature B-cells with sequences containing zero nucleotide mutations. In total, *n=*3,885 paired V_H_ and V_L_ sequences were processed from the *n=*20 Kymice.

### NGS Sequence Analysis of V_H_ Sequences from Bulk Sorted B Cells Derived from Kymice

Bone marrows from the femur and tibia of each Kymouse were processed into single-cell suspension as described above. From these *n=*20 bone marrow samples, seven were FAC sorted to recover CD19^+^ B220^+^ B cells into a single tube. The cells were lysed and RT-PCR was performed to amplify the V_H_ domain, followed by standard Illumina library generation, before sequencing on an Illumina MiSeq. The Change-O pipeline (Gupta et al. 2015) was used to process the sequences generated by the MiSeq sequencers. IgM sequences with zero mutations were selected for further analysis resulting in a total of 412,493 V_H_ sequences across the n = 7 Kymice (average 58,928 range 31,905 to 100,240).

### NGS Sequence Analysis of V_H_ Sequences Derived from Human Samples

Buffy coat samples were obtained from ten healthy individuals as described previously (Ghraichy et al, 2021). In the previous study, B-cells were FAC-sorted into naïve, marginal zone, plasma and switched memory cell populations. In the present study, we analysed IgM sequences from the naïve subset of B-cells with no mutations. There was a total of *n=*338,677 sequences (mean: 33,867 per human, range: 20,653 – 48,293).

### NGS Sequence Analysis of V_H_ Sequences Derived from C57BL/6 WT Mice

6,763,480 IgM V_H_ nucleotide sequences from naïve B cells of healthy unvaccinated C57BL/6 wild-type mice were downloaded from the Observed Antibody Space (OAS) (Kovaltsuk et al. 2018; Olsen, Boyles, and Deane 2022). Those sequences with any nucleotide mutations were removed, and the remaining sequences were down-sampled via stratified sampling aiming to preserve the original clonal structure in the complete dataset. 150,000 sequences with redundancy were randomly selected from each of the five C57BL/6 mice. Collapsing to unique (nucleotide) sequences resulted in a total of 268,285 sequences (mean: 53,657 sequences per mouse; range: 20,026-87,041).

### Clonal and diversity analysis of NGS sequence data

Clonotypes are defined as sequences with common IGHV and IGHJ genes and 90% or more amino acid identity across length-matched CDRH3s. Antibody sequences were assigned to clonotypes using the DefineClones module of Change-O (Gupta et al. 2015) under the amino acid model.

Shannon diversity (H) was calculated using the *stats.entropy* function within Python’s SciPy library.

The formula is as follows:

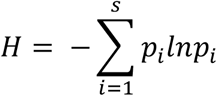

Where p_i_ is the proportion of sequences in the clonotype *i* of *s* clonotypes.

### Structural annotation

Sequences that IgBLAST identified as coding for a productive immunoglobulin were translated into amino acids and aligned to the IMGT antibody numbering scheme (Lefranc et al. 2003) using ANARCI (Dunbar and Deane 2016). IMGT CDR sequences were extracted and structurally annotated using the SAAB+ version 1.01 pipeline (Kovaltsuk et al. 2020). SAAB+ uses SCALOP (Wong et al. 2019) to assign each sequence’s CDRH1 and CDRH2 loops to a structural canonical class and uses FREAD (Choi and Deane 2010) to identify if the CDRH3 loop has a similar structure to any crystallographically-solved set of 4,544 CDRH3 structures (referred to as templates, downloaded from SAbDab (Dunbar et al. 2014; Schneider, Raybould, and Deane 2022) on 16^th^ February 2022).

To reduce dimensionality, templates are clustered with a 0.6Å RMSD cut-off, producing a set of 1,944 templates. In this set of templates, 41% of the antibody structures are of murine-origin and 37% are of human-origin (**Supplementary Figure 5**). The SAAB+ pipeline outputs for each sequence the canonical class of the CDRH1 and CDRH2 loops, and the Protein Data Bank (PDB) ID of the structure that contains the best matched CDRH3 structure for homology modelling from the 1,944 templates.

As a complementary approach, we also modelled representatives of all non-singleton clonotypes in each of the repertoires using a recent deep learning method, ABlooper (Abanades et al. 2022). ABlooper is competitive with AlphaFold2 on the canonical CDRs and shows better performance on the CDRH3 which was the target of our analysis. It is also more than 1000x faster than AlphaFold2 and was therefore suitable for our large-scale analysis.

For full structural modelling with ABlooper, a light chain must be supplied. As we did not know the cognate light chain for any of the heavy chains from bulk sequencing, all heavy chains were paired with a single light chain which was selected from the paired dataset. This light chain was the most commonly observed light chain in the Kymouse dataset. We selected a common light chain to standardise its effect on the prediction of the heavy chain CDRs, in the absence of knowledge of the true light chain.

#### Humanness scoring of V_H_ sequences

The human, Kymouse and C57BL/6 mouse V_H_ sequences were assigned a “humanness score” using the random forest regressors from Hu-mAb (Marks et al. 2021). Sequences were first IMGT numbered using ANARCI as above. While the C57BL/6 mouse and Kymouse sequences were mostly full length (IMGT positions 1-128), the human sequences were in most cases missing FWR1 (IMGT positions 1-26 in the IMGT CDR definition). As the human sequences were non-mutated, we considered it reasonable to simply fill in FWR1 according to the sequence found in the assigned germline. For the human and Kymouse sequences, the RF model trained on the IGHV gene assigned by IgBLAST were used for scoring, while for the murine sequences, all seven (IGHV1-7) RF models were used to score the sequence and the highest score was selected. We used the IGHV-specific classification thresholds reported in the Hu-mAb paper to annotate if a sequence was considered human or not (Marks et al. 2021).

#### Immunophenotyping of Intelliselect® Transgenic mice

Spleens, lymph nodes, and bone marrow from a further *n=*12 antigen-inexperienced Kymice were processed to single-cell suspensions and cryopreserved as described above. Fluorescently Activated Cell Sorting (FACS) of the bone marrow samples was performed using fluorescently conjugated antibodies against B220, IgM, IgD, IgG1, IgG2ab, IgG3, CD8, CD4, Ly-6G, CD11c and CD138 (BD Biosciences), CD19, F4/80, Sca-1 (BioLegend) and TACI (eBioscience). For the pooled spleen and lymph node samples the FACS panel consisted of B220, IgM, IgD, IgG1, IgG2ab, IgG3, CD8, CD4, Ly-6G&6C, CD11c and CD95 (BD Biosciences), CD19, F4/80, CD73, CD80, PD-L2 and GL7 (BioLegend). DRAQ7 (BioStatus) was used in all samples to distinguish live and dead cells. For flow cytometry, cells were thawed from frozen, resuspended in warm 10% FBS in RPMI buffer, filtered through 40 µm cell strainers and centrifuged at 400 × g for 10 minutes at 4 °C. Cells were resuspended in buffer and TrueStain FcX (BioLegend) was added for 10 minutes on ice. Single cell suspensions of bone marrow cells and pooled spleen and lymph node cells were stained with their respective staining cocktails for 30 minutes. All cells were spin washed and resuspended in buffer, filtered through a 30 µm cell strainer, followed by cell sorting on a 5-laser BD FACS Aria Fusion flow cytometer (Beckton Dickinson).

#### Analysis of Flow Cytometry Data from Intelliselect® Transgenic mice

The frequencies of the following cell types within total viable (DRAQ7) cells were determined using classical FACS gating: bone marrow plasma cells (CD138^+^TACI^+^/Sca-1^+^), spleen/lymph node memory B cells (B220^+^CD19^+^IgD^-^CD73^+^/CD80^+^/PDL2^+^) and spleen/lymph node germinal centre cells (B220^+^CD19^+^CD95^+^GL7^+^; data not shown). Cell populations were also analysed in an unbiased manner using unsupervised clustering algorithms. In brief, the raw .fcs files were imported into R (RStudio version 1.2.5033 with R version 4.0.0) using CytoExploreR (version 1.0.8) and the data were transformed to normalise marker intensities (logicle transform). For visualization, additional quantile scaling from 0-1 was performed, fixing values less than the 1^st^ percentile to 0.01 and values above the 99^th^ percentile to 0.99 to minimise the contribution of outliers to the scaling. Cell clustering was performed using the Leiden clustering algorithm (R package Monocle 2.16) and clusters were visualized in two-dimensional space using UMAP (R package uwot 0.1.10). Poorly resolved clusters were re-clustered, the subclusters manually merged to the first level clusters and annotated by cell type.

### Data availability

The sequence data is currently available in the Observed Antibody Space. Immunophenotyping data will be available upon publication.

### Code availability

The Python code used to analyse the data and generate the figures is available at https://github.com/oxpig/HumMus.

## Results

### Antigen naïve Kymice exhibit similar B cell sub-population frequencies

We characterised the B cell sub-populations within spleen and lymph node samples of 12 antigen-inexperienced Kymice using an 11-colour flow cytometry panel that incorporated a range of B cell lineage markers to identify both murine memory and germinal centre B cell populations. A canonical gating scheme organises B cells by their maturation status – from transitional B cells through naïve, non-switched and ultimately class-switched memory B cells. To look at the heterogeneity of the B cell subpopulations in more detail we incorporated unbiased Leiden clustering on the multi-parameter FACS data. Sorted cells separated into two large clusters, B-cells and non-B-cells (**Figure 1A**). As expected, within the B cell population immature isotypes (IgD and IgM) were enriched in naïve cells, whereas markers CD95 and GL7 were enriched in germinal centre cells (**Figure 1C**). The murine memory B cell population has been described to comprise five subpopulations defined by the progressive transition from naive-like to more memory-like cells and the surface markers CD80, CD73 and PD-L2 have previously been reported to enable their distinction (Tomayko et al. 2010). Using a low dimensional UMAP representation we observed distinct staining patterns of these markers in the memory B cell compartment and were able to distinguish between 12 major B cell populations, including transitional, naïve and activated as well as six distinct memory subsets. These were defined as (1) PD-L2^hi^, (2) CD73^hi^ CD80^hi^ PD-L2^low^, (3) CD80^low^, (4) PD-L2^hi^ CD80^hi^ CD73^low^, (5) CD80^hi^ and (6) CD73^hi^ **(Figure 1A)**. Germinal centre B cells formed a small and well separated cluster whose small frequency was not surprising given that these were antigen naïve animals. Based on the 3 memory markers (CD80, CD73 and PD-L2) the relative frequency of the total memory B cells was 6.60% ± 2.51%, and the frequency of CD95 and GL7 positive germinal centre cells was 0.18% ± 0.26%. The median expression profile of each subset is shown as a heatmap (**Figure 1C**). The antigen naïve B cell populations in un-immunised and non-infected Kymice are therefore normal and consistent between different Kymice (**Figure 1C right panel**).

**Figure 1:**
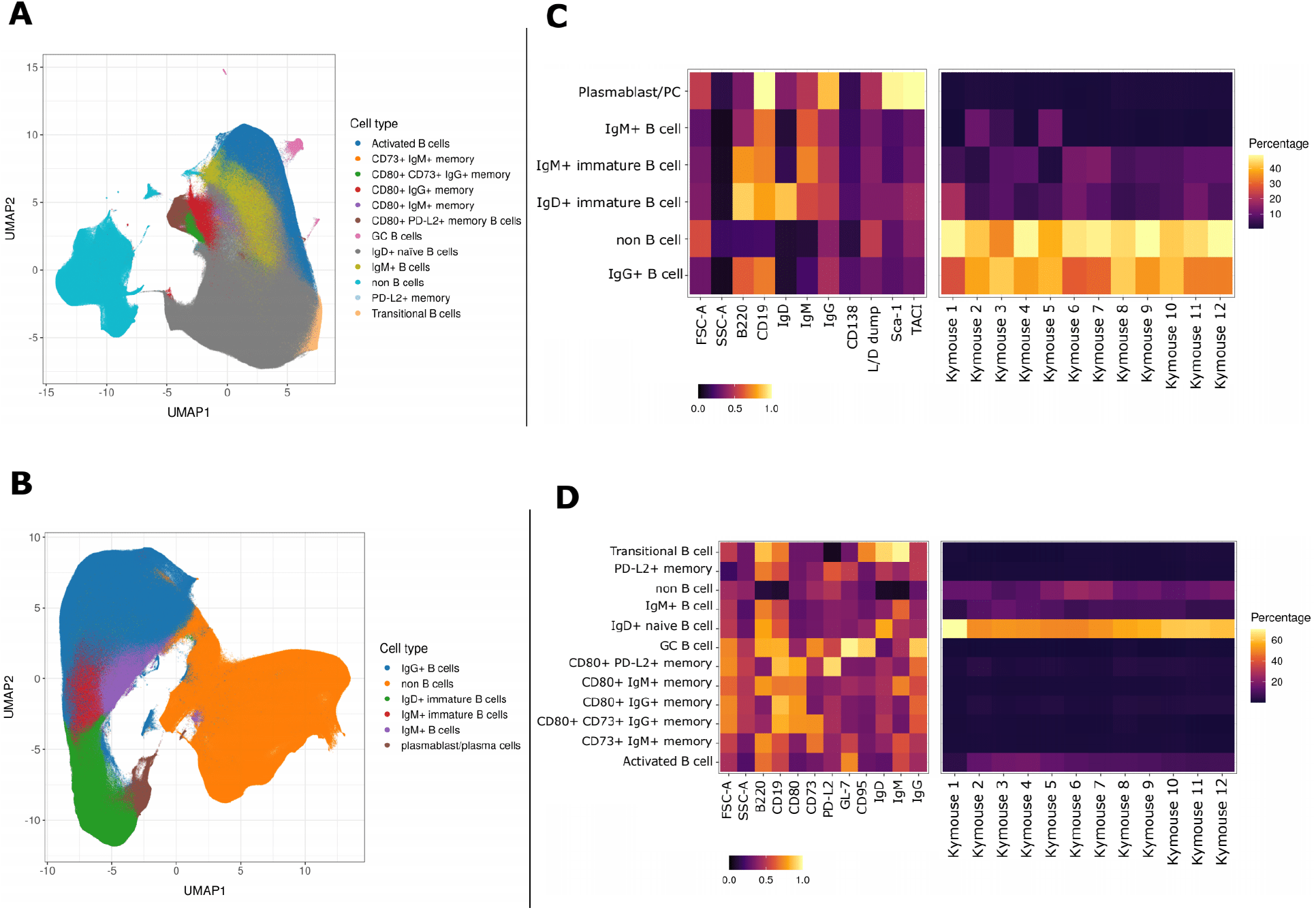
UMAP projections of sorted cell populations identified using unsupervised clustering from spleen and lymph nodes (A) or from bone marrow samples (B). UMAP projections show a clear separation between B-cells and non-B-cells for both sample types. The projections are coloured by the 12 resolved cell types in the spleen and lymph nodes (A) and the six resolved in the bone marrow samples (B). Normalised and scaled marker expression and frequencies showing mouse-to-mouse variation for each of the resolved cell types in the spleen and lymph nodes (C) or the bone marrow samples (D). The expression profiles are homogeneous across mice.

### B cells in the bone marrow are class switched with variable levels of surface BCR expression

To understand the B cell profile beyond spleen and lymph nodes we also profiled the bone marrow cells of mice. We characterized bone marrow samples using a 9-colour flow cytometry panel that incorporated a range of B cell lineage markers. The staining panel was designed to identify plasma cells as well as class switched B cell subsets in antigen inexperienced mice. As expected, we saw that the cells separated again into two large clusters, B-cells and non-B-cells **(Figure 1B**). The expression profiles of the subsets were again plotted as heatmaps showing the median expression profiles of each subset (**Figure 1D**). Within the B cell cluster, we identified several discrete subsets marked by the expression of different BCR isotypes. B cells were clustered into 5 distinct subpopulations, including immature IgD+ and IgM+ cells, mature IgM+ and IgG+ B cells and plasma cells. The markers TACI and Sca-1 were enriched in plasma cells as expected, whereas CD138, a common plasma cells marker did not show a high level of separation between the different cell types. Unsurprisingly, we saw the biggest separation between B cells and non B cells, and a continuum of B cell subtypes from IgD, through IgM, to IgG-expressing B cells as well as a discrete cluster identified as plasma cells. The frequencies of the plasma cells were low (0.90% ± 0.28%) in comparison to other B cell subtypes, perhaps not surprising given that these were antigen naïve animals.

### The Kymouse naïve antibody sequence repertoire is more human-like than murine-like

Using high-throughput sequencing we recovered 3,885 full-length paired V_H_ and V_L_ sequences and a further 451,655 full-length unpaired V_H_ sequences from naïve B cells extracted from the spleens and lymph nodes of 20 Kymice. In order to evaluate the humanness of the Kymouse naïve B cell sequence repertoire, we compared these sequences to equivalent datasets of 338,677 V_H_ sequences from human naive B cells, and 268,285 V_H_ sequences from C57BL/6 mice.

One of the most pronounced differences in heavy/light chain pairing between wild type mice and humans that has been described is the usage ratio of the Ig*κ* and Igλ chains in the BCRs of circulating B cells. Humans have an Ig*κ*/Igλ ratio of approximately 60:40 in serum and in mature B cells. Mice have an Ig*κ*/Igλ ratio of 95:5 in serum and 90:10 on B cells (McGuire and Vitetta 1981). We used the 3,885 full-length paired V_H_ and V_L_ sequences to calculate the Ig*κ*/Igλ ratio in Kymice and found a ratio of 51:49 (IQR: 55:45, 47:53), which is considerably closer to the human ratio than the mouse ratio.

We next used the unpaired V_H_ sequence data to determine the immunoglobulin heavy-chain gene usage frequencies in Kymice and compare the usage frequency of the IGHV, IGHD, and IGHJ germline genes to those in the human data. We used hierarchical clustering to compare the gene usage profiles of individual Kymice and humans, building phylogenetic trees to show the relationships between the individuals’ gene usage profiles. Although Kymice and humans are the same in the numbers of IGHV genes used the frequencies as determined by sequence abundance can differ. The hierarchical clustering of the IGHV genes showed that the Kymice and humans form nearly separate monophyletic clusters except for a single outlier human subject (Figure 2A). Most of the variation in IGHV gene usage is explained by the IGHV gene subgroup usage: clustering by this separates humans and Kymice without the outlier, with Kymice using a lower proportion of IGHV1 and IGHV2 genes compensated for by increased IGHV3, IGHV4 and IGHV6 usage (**Figure 2B**).

**Figure 2:**
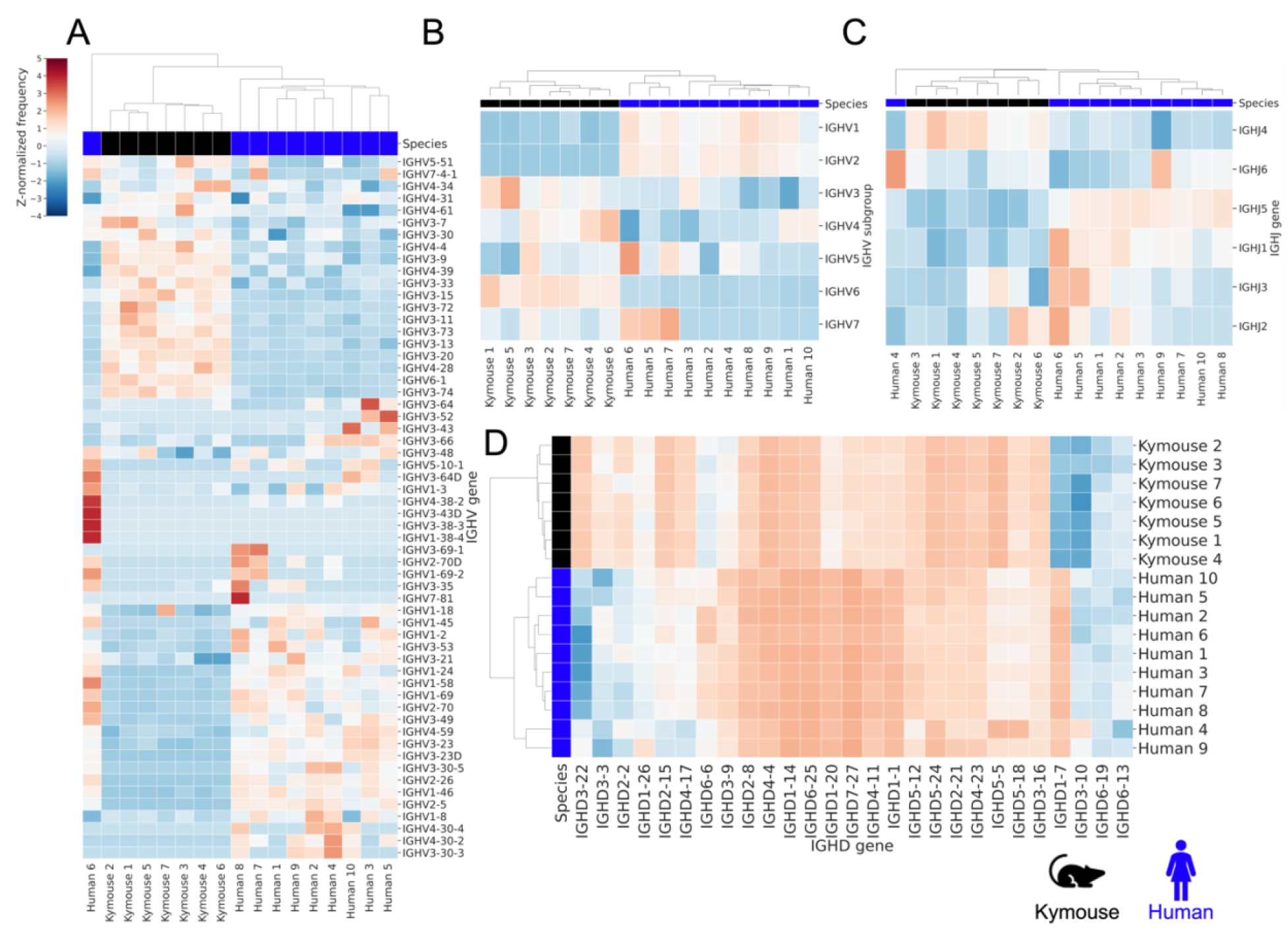
Gene usage clustermaps for A) IGHV genes, B) IGHV subgroups, C) IGHJ genes and D) IGHD genes. The IGHV clustermaps show a separation between human (blue) and Kymouse (black) repertoires, with lower usage of IGHV1 and IGHV2 in the Kymouse. Downstream, there are also differences in usage of IGHJ genes with a preference in the Kymouse repertoires for IGHJ4. The IGHD gene usage shows the clearest distinction.

The IGHJ gene usage profile is similar: Kymice and eight out of the ten humans form monophyletic clades with two outliers, including the outlier human subject from the IGHV gene usage clustering. On average, the Kymouse uses IGHJ4 more frequently than humans, and IGHJ5 and IGHJ1 less frequently (**Supplementary Figure 1**). Both the IGHV and IGHJ gene usage profiles of naïve Kymouse repertoires are more similar to one another than to any human repertoire. However, Kymice are closer to some of the human repertoires than others. This therefore indicates Kymice fall within the range of human IGHV and IGHJ usage frequencies.

The hierarchical clustering of the IGHD genes revealed that the Kymice formed a monophyletic sister group distinct to that of the humans (**Figure 2e**). As can be seen from the heatmap, the IGHD germline genes used by the Kymice (*e.g*. IGHD3-22, IGHD2-15) are infrequently used by humans and *vice versa*.

We compared the distribution of the CDRH3 lengths in each species’ naïve repertoire (**Figure 3A**). We calculated the length of the CDRH3 loop (under the IMGT antibody numbering scheme) from the Kymouse, human and C57BL/6 mouse heavy-chain sequences. This revealed that Kymice have an average CDRH3 length in between that of humans and mice, with a mean CDRH3 length of 14.25. In comparison, the C57BL/6 mouse dataset have a mean length of 12.39 amino acids, and the human dataset have a mean CDRH3 length of 16.56 amino acids. Kymouse CDRH3 loops are on average 2.36 aa shorter than humans (95% CI: 2.26, 2.48; p<0.001), while WT mice CDRH3 loops are on average 4.21 aa shorter than humans (95% CI: 4.12, 4.30; p<0.001).

**Figure 3:**
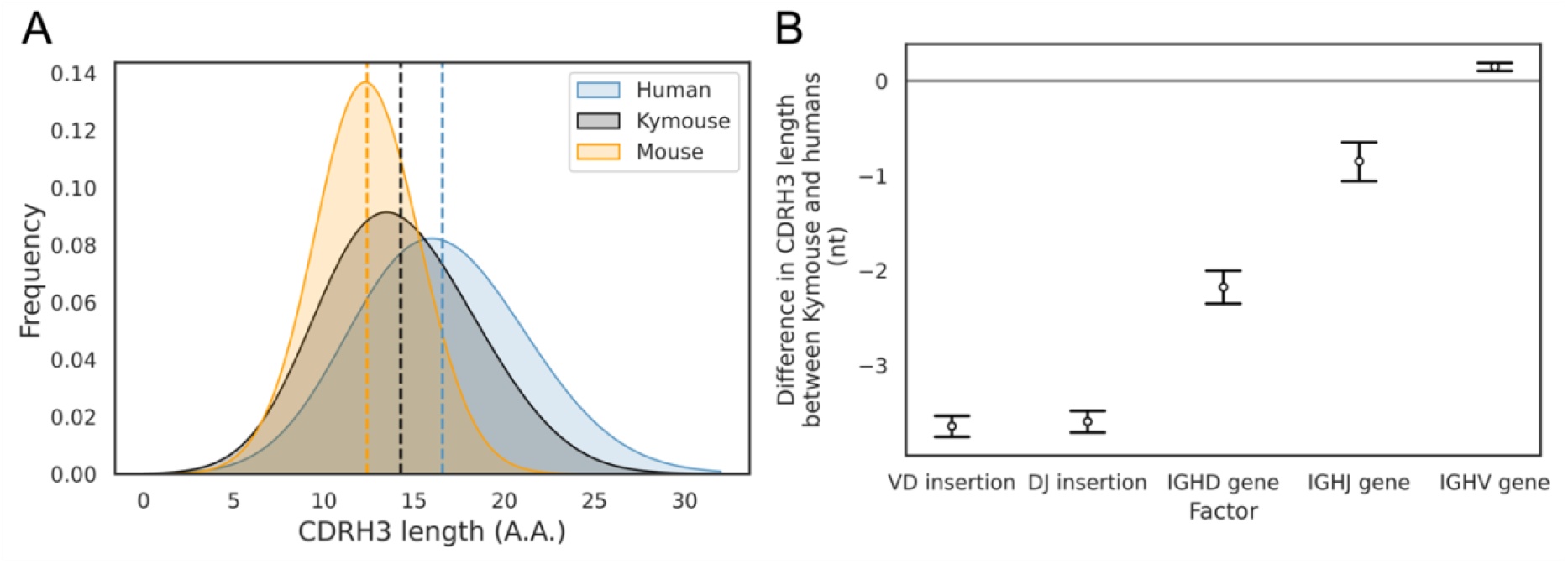
The CDRH3 length distribution among the mouse, Kymouse and human sequences. The Kymouse CDRH3 length distribution is intermediate between the mouse and human distributions (A). Plus or minus indicates the standard error of the mean. We used estimation statistics to reveal that the major factor leading to this reduction in CDRH3 length despite access to the same germline repertoire in Kymouse is the relative lack of VD and DJ insertions (B)(Supplementary Figure 3).

The length of a CDRH3 loop is determined by four main factors: (i) the choice of IGHV gene; (ii) the choice of IGHD gene; (iii) the choice of IGHJ gene; (iv) the number of nucleotides inserted or deleted in the V-D and D-J junctional regions during B cell development. In order to investigate further why the Kymouse tends to have shorter CDRH3 loops on average than humans, we considered each of these factors in turn. We first looked at whether the differential IGHV gene usage by Kymouse had a significant impact on the length of the CDRH3 loop. For each heavy-chain sequence in the human and Kymouse datasets we determined the number of nucleotides that the IGHV germline gene contributes to the CDRH3 loop. The results showed a statistically significant but small difference of 0.14 nt (95% CI: 0.12, 0.16; p<0.001) between humans and Kymouse. Therefore, it does not appear that the differential choice of IGHV between humans and Kymouse greatly affects the CDRH3 length. We next investigated the effect of the differential usage of IGHD genes between humans and Kymice on the length of the CDRH3 loop. This showed that the human IGHD germline genes used by the Kymouse are, on average, 2.34 nt shorter than humans (95% CI: 2.26, 2.41; p<0.001). We then compared the relative usage of the IGHJ genes of humans and Kymice (**Supplementary Figure 1**). Kymice tended to use IGHJ4 (47.3% in Kymice versus 44.2% in humans) and IGHJ6 (32.8% in Kymice versus 28.6% in humans) while using the other genes (IGHJ1, IGHJ2, IGHJ3, IGHJ5) slightly less frequently, in particular IGHJ5 (11.1% in Kymice versus 15.1% in humans). Estimation statistics revealed that the differential IGHJ gene usage between Kymouse and human resulted in a decrease in the CDRH3 length for Kymouse of 0.85 nt (95% CI: 0.77, 0.94; p<0.001).

Finally, we looked at the number of nucleotide insertions in the V-D and D-J junctions in the antibody heavy chains. The results showed that Kymice V-D junctions are on average 3.73 nucleotides shorter than humans (95% CI: 3.68, 3.78 p<0.001), with a mean insertion size of 3.35 nucleotides compared to 7.30 nucleotides for humans (**Supplementary Figure 2A**). Equally, the Kymice D-J junctions are on average 3.68 nucleotides shorter than humans (95% CI: 3.63, 3.73; p<0.001), with a mean insertion size of 2.91 nucleotides compared to 6.77 nucleotides for humans (**Supplementary Figure 2B**). Overall, the number of junctional nucleotides inserted in the Kymouse heavy chain is 7.33 fewer than in humans (95% CI: 7.25, 7.40; p<0.001), with an average of 7.55 in Kymice compared to 14.88 in humans. These results show that the main factors that give rise to the shorter CDRH3 lengths in Kymouse compared to humans are the reduced number of nucleotide insertions in the V-D and D-J junctional regions, and the differential usage of IGHD germline genes between the species (**Figure 3B**).

A correlate of shorter CDRH3 length is a reduction in the theoretical CDRH3 diversity obtainable, with each amino acid less in the CDRH3 leading to a further 20-fold reduction in the number of possible CDRH3s. We do not expect the difference in diversity to approach this magnitude because of the CDRH3 sequence patterns created by combinations of germline genes, however a reduction in the non-templated insertion of nucleotides could lead to less diverse repertoires in the Kymouse.

To examine the diversity of the repertoires, we considered how many unique CDRH3s or clonotypes can be found in a given repertoire, normalized by the total number of sequences observed. **Figure 4A** shows that the ratio of the number of unique CDRH3s to the number of sequences is comparable in Kymice and humans, and that both are considerably higher than in C57BL/6 mice. When clustering similar CDRH3s in combination with common IGHV and IGHJ genes (clonotypes), there are comparable unique clonotypes per sequence in mice, humans and the Kymouse (**Figure 4B**). We calculated the Shannon diversity of the CDRH3 and clone distributions, a measure which takes into account both richness and abundance and found comparable diversity across all repertoire types (**Figures 4C and 4D**). The diversity observed depends on CDRH3 length considered, with the mouse and Kymouse repertoires having greater diversity at shorter lengths (peak diversity at lengths 12 and 13 respectively) and the human repertoires having greater diversity at lengths greater than 18 amino acids (peak diversity at length 15) for both CDRH3s (**Figure 4E**) and clonotypes (**Figure 4F**). Given that junctional insertions are required to reach lengths of 23 or greater in the human and Kymouse data, this supports the hypothesis that the reduced junctional diversification in naïve Kymouse repertoires limits CDRH3 diversity at the longest lengths, however the greater diversity at shorter lengths may compensate. It is also clear that the Kymouse occupies a region in CDRH3 diversity and length between wild type mice and humans.

**Figure 4:**
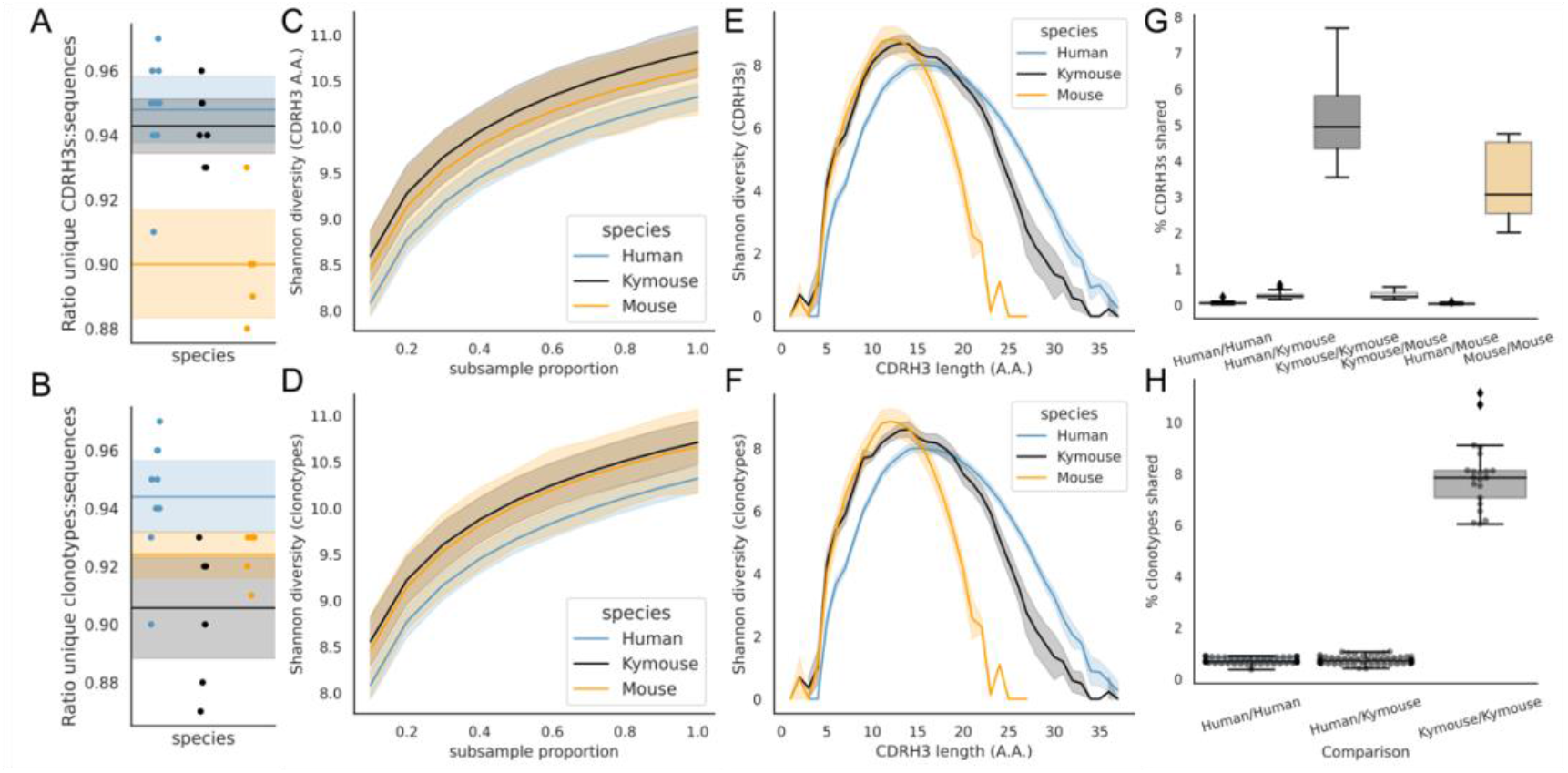
Examination of diversity of human, Kymouse and murine repertoires. The top rows pertain to exact CDRH3 (amino acids) and the bottom rows to clonotypes (same IGHV, IGHJ and greater than 90% amino acid identity across length-matched CDRH3s). At the level of CDRH3s, the Kymouse repertoires have more unique CDRH3s per sequence sampled (A) are more diversity in their usages (C), despite their limited VD and DJ insertion rates. Diversity is reduced relative to human sequences at longer CDRH3 lengths which in unmutated repertoires require VD/DJ insertions to reach (E). Kymouse repertoires show an opposite pattern in unique sampling rate and diversity when looking at clonotypes (B and D respectively) but still show reduced diversity versus humans at longer lengths. Overlap among CDRH3s (G) and clonotypes (H) between individuals is considerably higher in Kymice than humans, and more comparable to mice (G).

Finally, we considered the CDRH3 and clonotype overlap **(Figures 4G and 4H)**. The overlap among CDRH3s is highest between individual mice and individual Kymice. On average 5.1% of CDRH3s were shared between any two Kymice and 3.4% of CDRH3s shared between any two mice compared to 0.12% of CDRH3s shared between any two human subjects. Despite the greater CDRH3 diversity reported for individual Kymice, over 25 times more CDRH3s are shared between pairs of Kymice on average than between humans. Clonotype sharing was also higher between individual Kymice (average 7.92% vs 0.84% in humans).

### The Kymouse repertoires are structurally more human-like than mouse-like

Ultimately, genetic diversity is reflected in the structure of the resulting BCR and secreted antibody protein. Therefore, we compared the structural similarities of the BCRs via structural annotation of the CDRs. We first compared the CDRH1 and CDRH2s, which adopt a limited set of conformations known as canonical classes. For both CDRH1 and CDRH2 the Kymouse repertoires group separately from human repertoires, but group with human repertoires before the mouse repertoires, in their usage of canonical forms (**Supplementary Figure 3**). All canonical forms are strongly predicted by IGHV germline gene subgroup (**Supplementary Table 1**) especially as these sequences are non-mutated. Despite the effects of the differential IGHV usage observed in humans and Kymice, this does not make the Kymouse canonical class usage more similar to mice as the murine repertoires use different canonical classes than either the human or humanised repertoires. Interestingly, six of the nine murine IGHV germlines encode a single canonical form, H1-8-A, suggesting more limited structural diversity in murine CDRH1s.

We then performed structural comparisons of the shapes of the human, Kymouse and mouse CDRH3s. CDRH3s do not adopt canonical conformations, so we used two different approaches: firstly, structural annotation which consists of comparison to a CDRH3 structural database and annotation with a structural cluster ID, and secondly full CDRH3 modelling.

Of the 1,944 possible structural clusters in the CDRH3 structural database, 1,594 were observed in at least one repertoire. The majority (1,270) of these clusters were observed in all species. There was an observable difference among repertoires in the species origin of the structural clusters observed, i.e. each species was biased in the structural space it tended to use. The majority of structural clusters used in human and Kymouse repertoires were of human origin (57.5 and 55.0% respectively), while the majority of structural clusters used in mouse repertoires were of murine origin (64.0%) (**Supplementary Figure 4)**.

As described, the CDRH3 lengths in the human dataset were on average 2.36 aa longer than those in the Kymouse, and 4.21 aa longer than those in the C57BL/6 mice (**Figure 3A**). As we considered only CDRH3s between length 4 and 16 in this structural analysis, this difference was reduced to a difference of 0.45 amino acids between human and Kymouse repertoires (CI 0.43, 0.47) and 0.63 between human and mouse repertoires (CI 0.62, 0.65). The average CDRH3 lengths of the human templates in the FREAD database were longer than those in the murine templates (12.6 aa compared to 11.2 aa respectively) (**Supplementary Figure 5**). We checked that the observed differences in structural template usage by humans, Kymice, and C57BL/6 mice were not just reflecting differences in the availability of templates at the difference CDRH3 lengths. Therefore, we stratified the FREAD database by CDRH3 length and ran the datasets against each CDRH3 length separately. If the preferential usage of human-specific templates by humans and humanized mice, and vice-versa for mice, is simply a result of random sampling from the FREAD database, we expect the ratio of the proportion of human templates in human repertoires to the proportion of human templates in the FREAD database to be approximately 1 across all CDRH3 lengths, and below 1 across all CDRH3 lengths for mice. Instead, we see a significant enrichment (at the 1% level) of species-specific templates at multiple CDRH3 lengths.

The human repertoires are more structurally variable than are the humanized murine or murine repertoires (**Figures 5A, B and C**). The murine repertoires are the least variable (there is the smallest range of distances between any two given subjects for mice, and on average the smallest average distance between pairs of subjects). All the Kymouse repertoires were structurally closer (Euclidean distance in Z-normalized CDRH3 structural cluster usage) to any given human repertoire than to any C57BL/6 mouse repertoire **(Figures 5A, B and C**). With no correction for sample size or for CDRH3 length distribution, the humanized murine and human repertoires form a monophyletic cluster that is sister group to the murine repertoires (**Figure 5D**).

**Figure 5:**
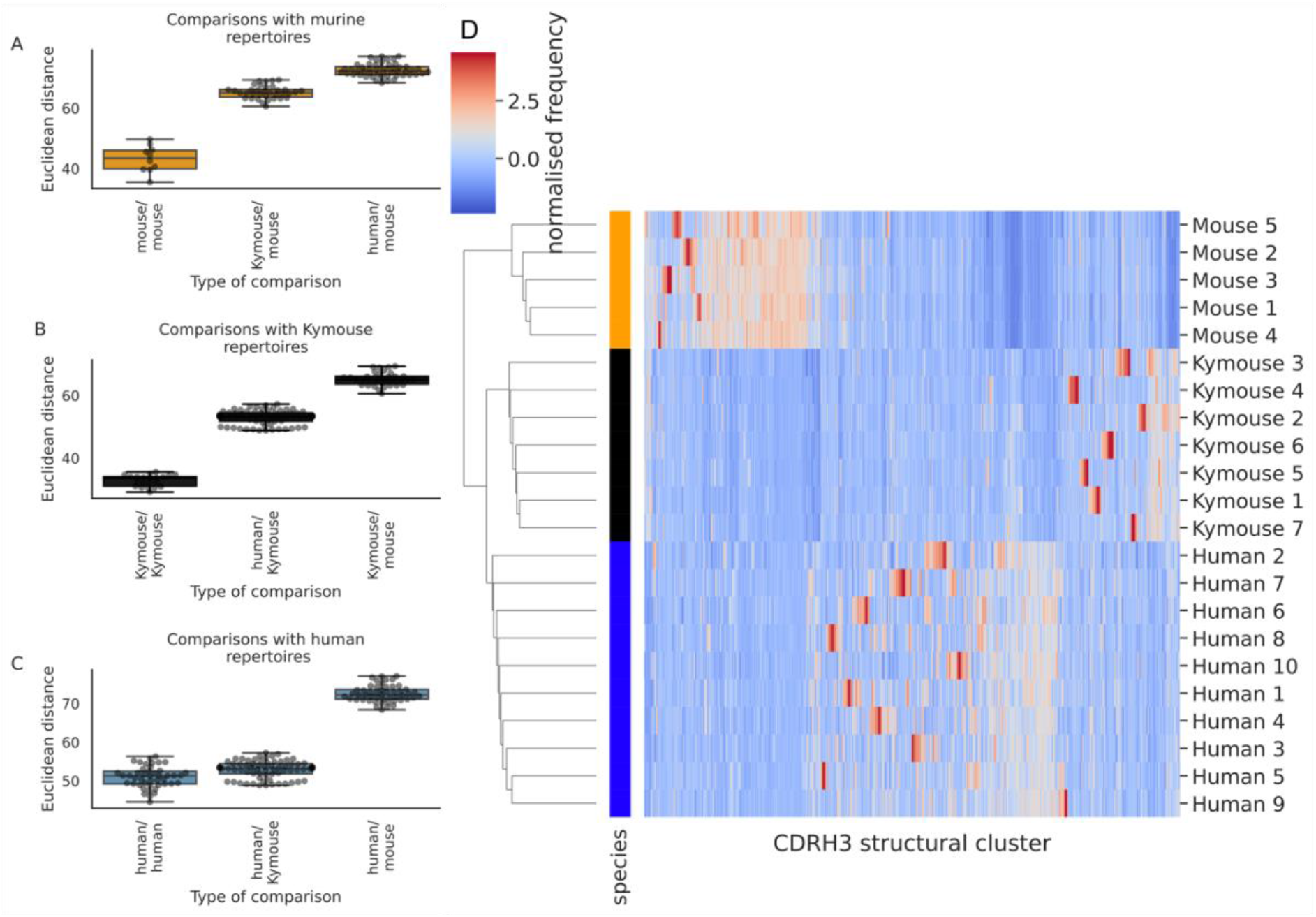
Distance between CDRH3 structural cluster usage profiles is measured with Euclidean distance in Z-normalized proportions. This is calculated pairwise between subjects and these distances are clustered hierarchically. Figures A through C show these pairwise distances stratified by the type of comparison. The leftmost box shows the range in distance between individuals of the same species. For the mouse and Kymouse repertoires, this range is smaller than the range in distances for any other species, meaning that they cluster monophyletically (D). Human repertoires have less self-similar CDRH3 structural cluster usages with ranges overlapping with Kymouse repertoires. In the hierarchical clustering solution with these pairwise distances that is shown in D, the human and Kymouse repertoires form a monophyletic clade separately from the murine repertoires.

Labelling cluster members as “murine” or “human-like”, the adjusted Rand index of this clustering is 1.0 (perfect correspondence). We calculated Rand indices for 100 subsamples with equal numbers of each CDRH3 length to control for any length effect; in all subsamples, the human and humanized repertoires could be clustered separately from the murine repertoires resulting in an adjusted Rand index of 1.0. Further, in all 10 subsamples, at least one human repertoire was more structurally similar to a given humanized murine repertoire than to at least one other human repertoire (between 5 and 9 of 10 subjects per subsample).

In conclusion, even when adjusting for sample size and differing CDRH3 length distribution, the repertoires of Kymouse are structurally more similar to human repertoires than they are to murine repertoires according to the homology-based structural annotation technique.

The species bias we observed in structural cluster usage, which is assigned via sequence similarity to a structural template, meant that a template-free modelling procedure might lead to different results. We modelled non-singleton clonotype representatives from humans, Kymice and mice using ABlooper, then compared the resultant CDRH3s via C_α_ RMSD. Analogously to clustering of the CDRH3 template database, we performed greedy clustering with a 0.6Å cut-off. This clustered the 43,378 models with 41,397 unique CDRH3s into 6,546 clusters. The modelled human CDRH3s were on average longer than Kymouse CDRH3s by 0.99 amino acids (CI: 0.94, 1.05) and longer than mouse CDRH3s by 1.23 amino acids (CI: 1.17, 1.29). We observed a slightly different clustering by usage than via the homology approach (**Supplementary Figure 6A**), with monophyly of Kymice, and Kymice and humans, but not of mice. The distribution of intersubject differences was such that the human/Kymouse and human/human comparison distributions are largely overlapping. Indeed, the differences between all distributions were less extreme with overlap between most of the distributions **(Supplementary Figure 6B**). The monophyly of Kymice and humans versus mice is observed when subsampling in order to equalize the CDRH3 length distribution, i.e., it is not driven by differences in CDRH3 length. The deep learning-based modelling approach supports the earlier finding that Kymice are more structurally similar to humans than mice, and vice-versa; the extent of this similarity is greater than observed with the homology modelling approach.

In conclusion, the sequence differences in the repertoires which were described in the germline and diversity sections do impact upon the repertoire of CDR structures which are observed in the Kymouse. The distribution of canonical forms in CDRH1 and CDRH2 is distinguishable, but usage is human-like. In the CDRH3, both structural annotation and full structure prediction indicate that the naïve Kymouse CDRH3 structural space is human-like, indicating that Kymice repertoires offer comparable structural starting points for the production of antigen-specific antibodies.

### A state-of-the art humanization tool scores Kymouse sequences as fully human

We next tested whether the Kymouse sequences are considered human by a state-of-the-art humanization tool. We used the random forest classifiers within Hu-mAb to score heavy chain amino acid sequence humanness. 100% of human and Kymouse heavy chain sequences were classified as human, with the maximum humanness score assigned to 99.1% of human sequences and 98.3% of Kymouse sequences. All sequences produced scores in the “Positive (High Score)” category which had minimal anti-drug antibody events reported among therapeutic antibodies. No murine sequences were classified as human.

## Discussion

The phenotypic diversity of B cells in the spleens, lymph nodes and bone marrows of the Kymouse, determined by immunophenotyping panels, showed the main immunologically relevant B cell subpopulations could be identified at appropriate cell frequencies, consistent with the Kymouse being fully competent for B cell development and capable of a complete humoral immune response. As expected, the baseline levels of the immune relevant subsets, i.e. memory B cells and germinal centre B cells in the spleen and lymph nodes as well as the plasma cells in bone marrow were low reflecting the lack of immune exposure beyond commensal and environmental antigens during mouse husbandry.

B-cells recognise antigens via the B-cell receptor (BCR) and an individual is able to generate a set of high-affinity BCRs to any given antigen due to the exceptional genetic and structural diversity of B-cell receptor binding sites created by recombination of germline genes, junctional diversification and somatic hypermutation. While somatic hypermutation plays a key role in the development of mature high affinity antibodies, the breadth of an immune response to an antigen is limited at first by the diversity produced by recombination and non-template additions of the non-somatically hypermutated, germline encoded heavy chain immunoglobulin genes and their pairing with similarly rearranged immunoglobulin light chain genes that together comprise the BCRs in the naïve B-cells.

Paired V_H_/V_L_ sequencing of the naïve Kymouse BCR repertoire revealed a near 50:50 Ig*κ*/Igλ ratio which more closely approximates the human Ig*κ*/Igλ ratio of 60:40 than the murine Ig*κ*/Igλ ratio of 95:5. Sequence analysis of immunoglobulin heavy chains showed that naïve Kymouse BCRs have a CDRH3 length distribution that is intermediate between human and mouse repertoires (on average ∼14 aa versus ∼16 aa for human repertoires and ∼12 aa for mice). We compared the IGHV, IGHJ and IGHD gene usage profiles of human and Kymouse repertoires and showed that the IGHV and IGHJ gene usage profiles in Kymice are distinct from human profiles but human-like, while the IGHD gene usage profiles are sufficiently different that Kymouse and human repertoires are distinct. This contrasts with the previous NGS characterization of the OmniRat in which both IGHV and IGHD gene usage was distinct from humans (Joyce, Burton, and Briney 2020). It is an ongoing concern that different germline gene distributions may affect how representative transgenic models are of immune responses to germline-targeting immunogens.

We then examined the extent to which the differing germline gene usage distributions contribute to the shorter CDRH3s observed in Kymice. The Kymouse repertoires display a preference for shorter IGHD genes: this was also observed in the NGS characterization of Omni-Rat BCR repertoires suggesting that a preference for shorter IGHD genes may be common across transgenic rodent platforms (Joyce, Burton, and Briney 2020). While the differences in germline gene usage distributions do appear to contribute to the differing CDRH3 length distributions, a greater proportion of the effect is ascribable to fewer nucleotide insertions at both the V(D) and (D)J junctions during junctional diversification in the Kymouse with on average 7.33 nt fewer inserted in Kymouse over the two junctions. This reduced junctional diversification in the Kymouse leads to lower diversity in longer CDRH3s and greater clonotype overlap between individual Kymice than between individual humans.

Changes in CDRH3 structural cluster usage have been previously observed along the B cell development axis (Kovaltsuk et al. 2020) and allow comparison of repertoires derived from different species. Despite these genetic differences, modelled structural comparison of the human and humanized repertoires to the murine repertoires, by annotating the V_H_ sequences with predicted CDRH3 structural template clusters showed murine repertoires use mostly CDRH3 structural clusters that have been identified from murine antibodies, while human and Kymouse repertoires use CDRH3 structural clusters identified from a more even distribution of species of which more than 50% were identified from human antibodies. Further, grouping of the exact distribution of CDRH3 structural clusters reveals that Kymouse structural cluster usage is closer to human usage than murine usage, accounting for CDRH3 length differences. When modelling the CDRH3s (as opposed to performing an approximation via structural annotation), part of the “structural distance” between the human, mouse and Kymouse repertoires disappeared suggesting part of the signal observed in homology modelling is due to different sequences with similar predicted shapes. This suggests that despite the observed differences at the sequence level, the CDRH3 structural shapes adopted by the BCRs are within the distribution of observed human shapes.

Finally, using the Hu-mAb humanness classifiers, all Kymouse and human sequences are classified as human, meaning that naïve sequences isolated from the Kymouse are predicted to have similar immunogenicity in humans to sequences isolated from humans themselves.

In conclusion, although naïve BCR repertoires of the Kymouse have key distinctions from human repertoires at the sequence level they are comparable to the human repertoires in terms of CDRH3 structural usage. A number of studies have shown how the Kymouse is able to elicit equivalent antibodies to those found in humans exposed to the same antigen (Sok et al. 2016; Scally et al. 2017; McLeod et al. 2019; Oyen et al. 2020). This suggests that the engagement of BCRs on naïve B cells is authentic, and that the structural templates available for antigen binding are indeed human-like as we show here.

