## Supplementary Information for "Characterisation of the immune repertoire of a humanised transgenic mouse through immunophenotyping and high-throughput sequencing"

Richardson E[1,2*], Binter Š[1*], Kosmac M[1], Ghraichy M[3,4], von Niederhausern V[3,4], Kovaltsuk A[2], Galson J[5], Trück J[3,4], Kelly DF[6], Deane CM[2], Kellam P[1, 7], Watson SJ[1]

[1] Kymab, a Sanofi Company, Babraham Research Campus, Cambridge, UK

[2] Department of Statistics, University of Oxford, Oxford, UK

[3] Division of Immunology, University Children's Hospital, University of Zurich, Zurich, Switzerland

[4] Children's Research Center, University of Zurich, Zurich, Switzerland

[5] Alchemab Therapeutics Ltd, Kings Cross, London, UK

[6] Department of Paediatrics, University of Oxford, Oxford, UK

[7] Department of Infectious Diseases, Faculty of Medicine, Imperial College London, UK

[*] Contributed equally to the study

| Species | IGHV subgroup | Dominant H1 canonical form | Dominant H2 canonical form | % in dominant H1 canonical form | % in dominant H2 canonical form |
|---|---|---|---|---|---|
| Human | IGHV1 | H1-8-A | H2-8-A | 60.5 | 100.0 |
| Human | IGHV2 | H1-10-A | H2-7-A | 100.0 | 100.0 |
| Human | IGHV3 | H1-8-A | H2-8-B | 100.0 | 86.7 |
| Human | IGHV4 | H1-8-A | H2-7-A | 63.7 | 100.0 |
| Human | IGHV5 | H1-8-C | H2-8-A | 100.0 | 100.0 |
| Human | IGHV6 | H1-10-B | None | 100.0 | 100.0 |
| Human | IGHV7 | H1-8-C | H2-8-A | 99.9 | 100.0 |
| Kymouse | IGHV1 | H1-8-C | H2-8-A | 66.3 | 100.0 |
| Kymouse | IGHV2 | H1-10-A | H2-7-A | 100.0 | 100.0 |
| Kymouse | IGHV3 | H1-8-A | H2-8-B | 100.0 | 78.9 |
| Kymouse | IGHV4 | H1-8-A | H2-7-A | 46.2 | 100.0 |
| Kymouse | IGHV5 | H1-8-C | H2-8-A | 66.3 | 100.0 |
| Kymouse | IGHV6 | H1-10-B | None | 100.0 | 100.0 |
| Kymouse | IGHV7 | H1-8-C | H2-8-A | 100.0 | 100.0 |
| Mouse | IGHV1 | H1-8-A | H2-8-A | 49.9 | 96.7 |
| Mouse | IGHV2 | H1-8-A | H2-7-A | 99.9 | 100.0 |
| Mouse | IGHV3 | H1-9-A | H2-7-A | 90.2 | 100.0 |
| Mouse | IGHV4 | H1-8-A | H2-8-B | 81.4 | 100.0 |
| Mouse | IGHV5 | H1-8-A | H2-8-B | 98.1 | 100.0 |
| Mouse | IGHV6 | H1-8-A | H2-10-A | 99.9 | 100.0 |
| Mouse | IGHV7 | H1-8-A | H2-10-A | 100.0 | 100.0 |
| Mouse | IGHV8 | H1-10-A | H2-7-A | 100.0 | 94.3 |
| Mouse | IGHV9 | H1-8-A | H2-8-A | 100.0 | 100.0 |

**Supplementary Table 1**: dominant canonical forms per IGHV subgroup. Predicting the canonical form based solely on the dominant form observed for the IGHV subgroup would result in accuracy between 46.2 and 100%. Differences can be seen between the Kymouse and human in IGHV1, in which the majority of human sequences are H1-8-A vs. H1-8-C in the Kymouse.

| CDRH3 length (A.A.) | Number of unique CDRH3s | Number of clusters with 2Å cut-off | Number of clusters with 1Å cut-off | Number of clusters with 0.6Å cut-off |
|---|---|---|---|---|
| 4 | 25 | 2 | 2 | 5 |
| 5 | 43 | 2 | 2 | 6 |
| 6 | 129 | 2 | 5 | 12 |
| 7 | 309 | 2 | 7 | 26 |

| | | | | |
|---|---|---|---|---|
| 8 | 640 | 4 | 13 | 60 |
| 9 | 1304 | 4 | 24 | 115 |
| 10 | 4348 | 6 | 30 | 175 |
| 11 | 6373 | 6 | 37 | 264 |
| 12 | 6799 | 6 | 69 | 448 |
| 13 | 6828 | 12 | 111 | 894 |
| 14 | 6040 | 16 | 188 | 1275 |
| 15 | 4790 | 19 | 220 | 1460 |
| 16 | 3769 | 23 | 289 | 1806 |
| **Total** | **41,397** | **104** | **997** | **6546** |
| Metric | | Metric at 2Å | Metric at 1Å | Metric at 0.6Å |
| Kymouse and humans monophyletic | - | True | True | True |

**Supplementary Table 2:** information about the number of CDRH3 structural clusters produced with different thresholds under the select greedy clustering algorithm. 0.6Å was the threshold selected by Kovaltsuk and colleagues (Kovaltsuk et al, 2020) in the original SAAB+ application.

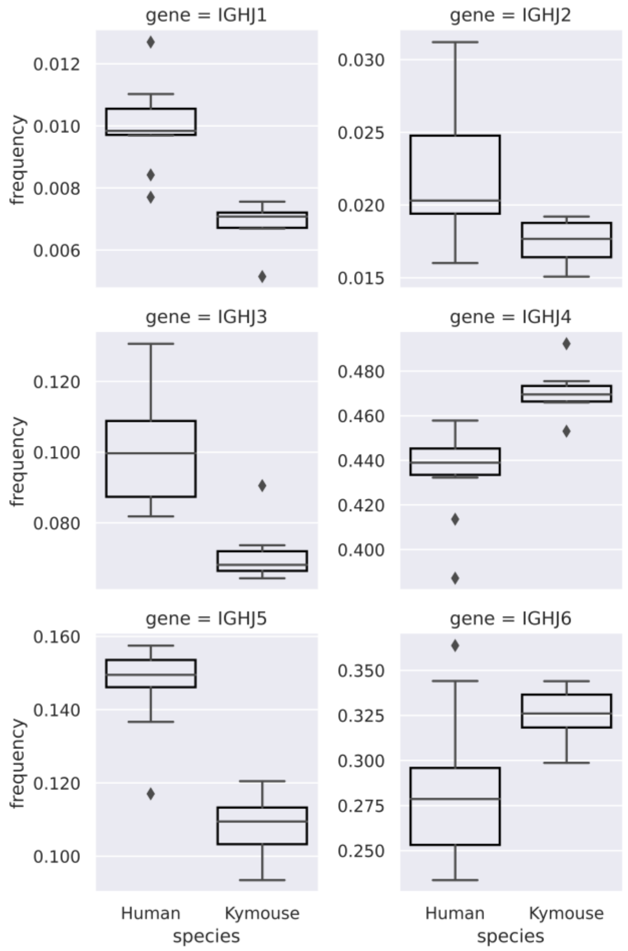


**Supplementary Figure 1:** comparison of IGHJ usage between naïve, human and Kymouse IGHM BCR repertoires. Kymouse repertoires disproportionately use IGHJ4 and IGHJ6 versus human repertoires, and use less IGHJ1, IGHJ2, IGHJ3 and IGHJ5.

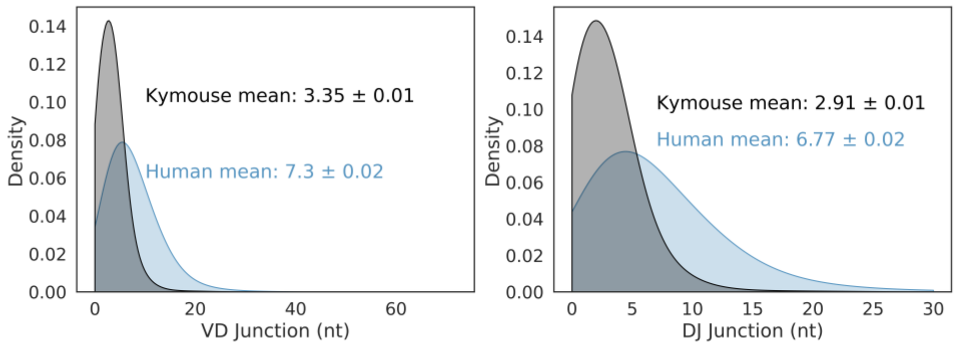


**Supplementary Figure 2**: VD and DJ insertion length distributions in the Kymouse (grey) versus human (blue) repertoires. We considered only sequences where a satisfactory IGHD germline alignment was achieved.

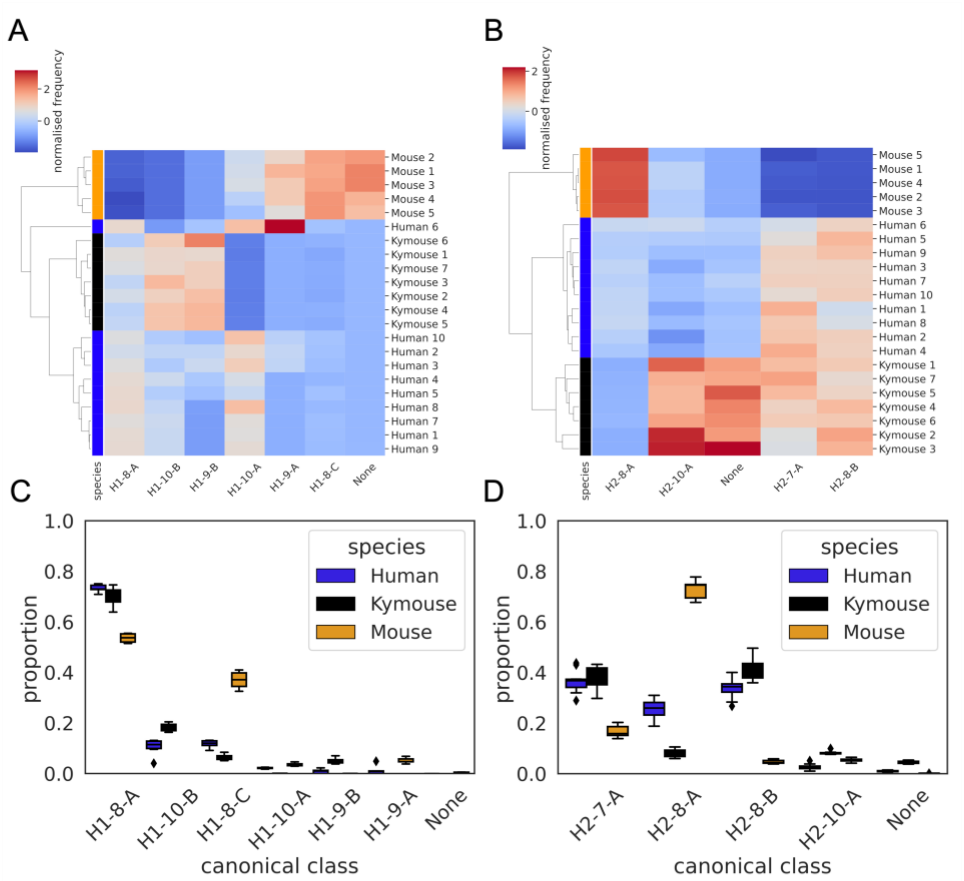


**Supplementary Figure 3:** CDRH1 (A and C) and CDRH2 (B and D) canonical class usage clusters Kymice and humans separately from mice. The key differences between humans and Kymice and mice in CDRH1 canonical forms are the greater usage of H1-9-A and H1-8-C in mice verus H1-10-B and H1-9-B in humans and Kymice. Between humans and Kymice, the Kymouse repertoires have lower usage of H1-10-A and greater usage of H1-10-B than humans. Focussing on the CDRH2 canonical forms, the largest difference between mice and humans/Kymice is in the greater usage of H2-8-A and lower usage of H2-8-B and H2-7-A. The Kymouse repertoires use significantly less H2-8-A than the human repertoires.

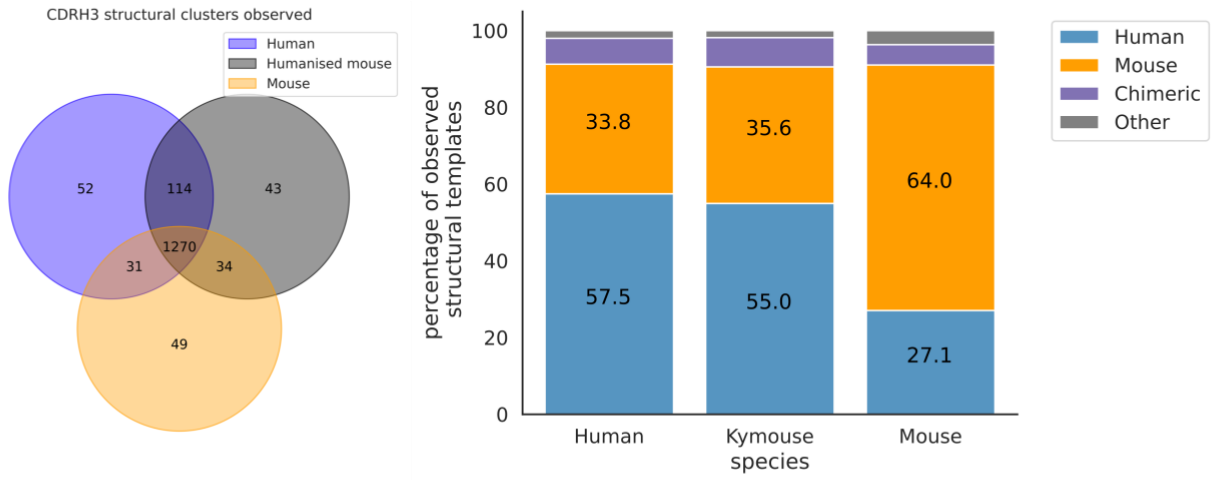

**Supplementary Figure 4:** the majority of CDRH3 structural clusters are observed across all three repertoire types. Focussing on the origin of the antibody representing the structural cluster, 57.5% of templates used in the human repertoires and 55% in the Kymouse repertoires are of human origin. The majority of templates observed in the mouse repertoires are of murine-origin.

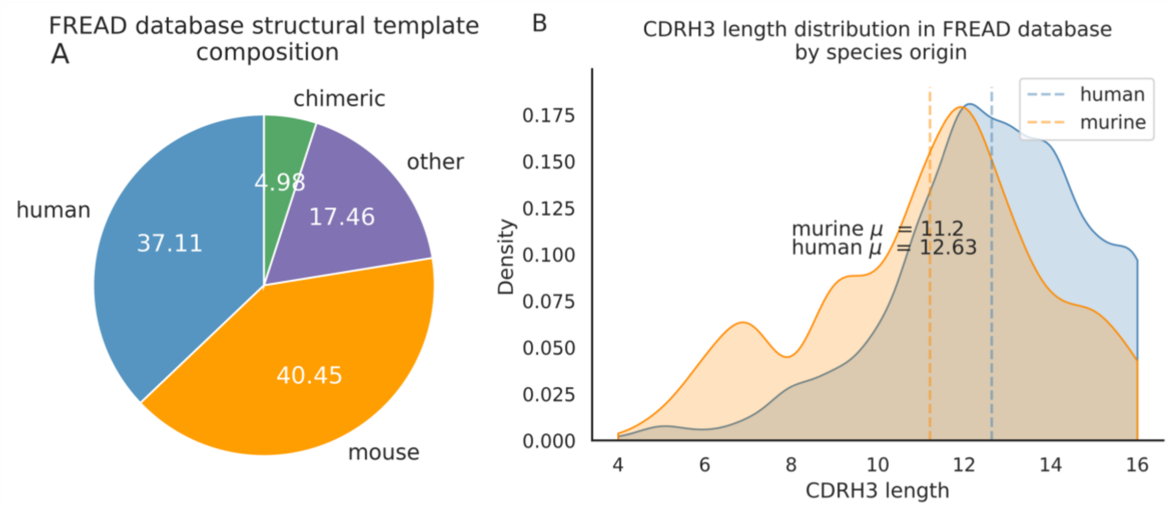


**Supplementary Figure 5:** around 40.5% of templates in the FREAD database are labelled as of murine-origin vs. 37.1% of human templates. There are proportionally more human templates at CDRH3 lengths of longer than 12 amino acids.

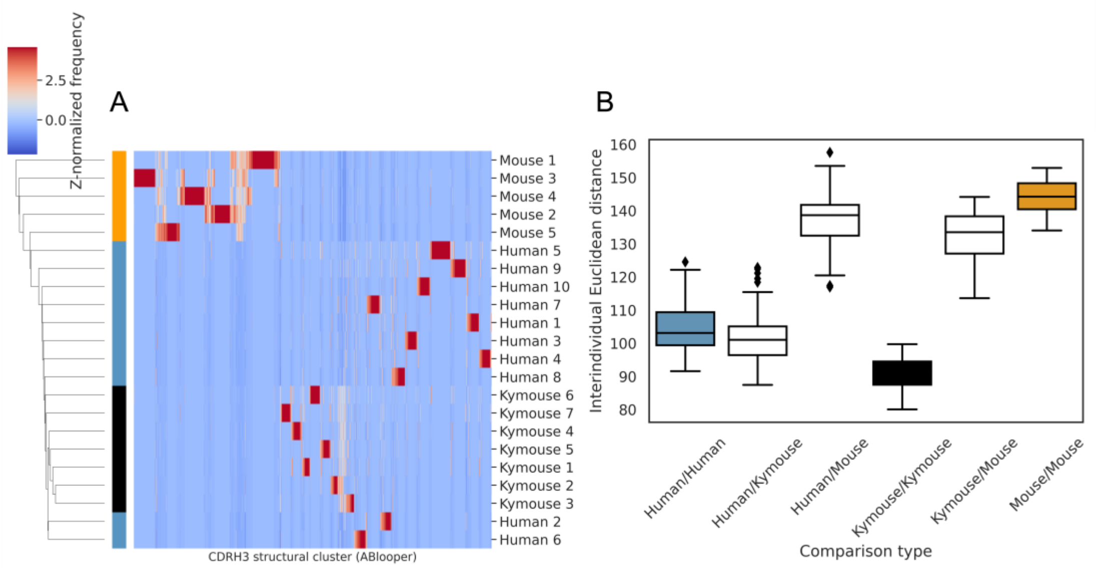


**Supplementary Figure 6**: A shows the clustermap of Z-normalized usages of CDRH3 structural clusters which were derived from greedy clustering with a 0.6Å cut-off of CDRH3 Cα RMSDs calculated between models built by ABlooper. The humans and Kymice form a monophyletic cluster, which the C57BL/6 mouse repertoires do not. Kymice and humans form monophyletic clades. The distribution of distances shown in figure B reveals that the Kymouse repertoires are the least variable. Similarly to the SAAB+ CDRH3 structural cluster usage comparison, the ranges of human/human and human/Kymouse intersubject distances are overlapping however the extent of overlap is greater than observed with SAAB+. There is also overlap between the distances observed between humans and Kymice with mice, and between individual mice.
